# Arduino-based lab equipment: building a multipurpose pressure transducer device

**DOI:** 10.1101/2020.09.13.295097

**Authors:** Stefano Di Domenico

## Abstract

**Background:** Lab equipments could be expensive and their cost can be unsustainable for scientist with limited financial resources. In order to overcome these impediments and to improve our experimental studies on liver resection in rat, a multipurpose pressure measurement device was project and realized using low cost components.

**Materials and Methods:** The device is based on an Arduino board, an easy-to-use and open-source microcontroller, that receives analog inputs from an instrumental amplifier connected to a disposable pressure traducer. The analogic inputs are converted to digital values and an LCD can visualize the pressure values calculated from the digital inputs. Programs has been written using C++ within the Arduino IDE, while the pressure data has been recorded on PC using data-logging freeware. Calibration has been performed using a water column as standard. Measure agreement was studied by comparing this device with a water column and with a standard clinical monitoring system.

**Results:** The final device it is handy and portable, it costs less the 20 euros and both the hardware and the software are easy modifiable. Calibration procedures resulted in reliable measurements as showed by the Bland-Altman measure agreement analysis.

**Discussion and conclusion:** The final device meets the goals of the original project and offers pressure measurements that are enough accurate for our studies on liver regeneration.

In a wider contest the present article shows the potentiality of the open-source-hardware movement to increase the opportunities for scientists and educators.

## Introduction

Conventional lab equipment can be expensive, limiting their availability and lessening the chance of testing new ideas and start new projects.

However, many open-source micro-controllers and sensors are available at low cost for educational and recreational purposes: they do not require an academic-level knowledge of electronics or informatics, and they have been employed by researchers to set up many type of lab instruments [1–4].

Inspired by these experiences, an Arduino-based pressure measurement device was developed, in order to measure and monitor the portal and caval pressure during our research protocol of extensive liver resection in rat [5].

## Materials & Methods

### Device design

A portable device was designed in order to measure, display and eventually store data of intraluminal blood pressure ranging from 0 to 40 mmHg.

This instrument was built using components that have been identified, tested and assembled after considering their availability and cost:

1. an open-source microcontroller board
2. a pressure transducer commonly used in the operative room and in the intensive care unit for invasive blood pressure monitoring
3. an instrument amplifier based on the OP90 integrated circuit
4. a LCD
5. a main switch, a push button, and few passive electronic components such as resistors, capacitors, trimmers, and wires
6. an electrical junction box

### The microcontroller

Arduino board is a family of open-source, low-cost, integrated circuits that contain an Atmel AVR processor, memory and analog and digital input and output pins. These boards can be programmed in C++ within the Arduino Integrated Development Environmental (IDE) in order to acquire inputs from a variety of switches and sensors and to control lights, motors, and other physical outputs [6]. The Arduino boards are open source: their schematics, hardware, design files, and source code are released under a Creative Commons Share Alike license and they are freely available for viewing and modification. Since anyone is free to modify the hardware design and to produce their own version, several low-cost clones are available, and one of them was selected for this project (DccEle – DCcduino UNO).

This board is based on the ATMEGA328P (16MHz), it offers 14 digital input/output pins, and 6 analog input pins working with 10 bits of resolution, leading to detect 1024 different values from 0 to 5 volts. The board can be powered via USB connection or with an external power supply from 6 to 20 volts (V): in this project a supplemental internal power supply has been design using a common 9 V battery.

Digital inputs and outputs can be recorded on a PC using CoolTerm, a freeware serial port terminal application that establishes a connection between computer and external devices in order to exchange and analyze data. It is freely available at http://freeware.the-meiers.org.

### Pressure Transducer

TruWave is a disposable pressure transducer distributed by Edwards Lifesciences® that communicates blood pressure information from a pressure catheter to a patient monitoring system [7]. It is widely used in operative rooms and intensive care units. Despite its main specifications are available on the producer website, the construction details are patent-protected, and some assumptions have been made and tested in order to use it for our purposes. It was assumed that TruWave is based on a Wheatstone bridge as many others pressure sensors, and the red and black wires that compose its connector line are the power wires while the others are the sensor outputs. Based on that hypothesis the sensor was supplied by 5 V DC and the output wires were checked using a multimeter by connecting the sensor to a water column: the output voltage of two wires (green and white) was proportional to the pressure, while the yellow one output showed a stable voltage value.

### Instrumental amplifier

The sensor output voltage is in μV (10^^−6^ volts) and it must be amplified in mV (10^^−3^ volts) in order to be clearly detectable by the microcontroller-board, since its resolution is around 4,9 mV (5V / 1024). For this purpose a simple amplifier was build, based on the OP90 integrated circuit: following the instructions contained in its data-sheet [8], a single-supply instrumentation amplifier was assembled, checking different sets of resistors to achieve the desired pressure range amplification.

### Visual Output

An LCD display was used to visualize the pressure values and the battery charge: it is able to display 16 characters arranged in 2 rows, and it can be controlled by the microcontroller board with an I2C module using only two digital pins.

### Battery charge monitor

In order to monitor the charge of the internal 9 V battery, a symmetric voltage divider was connected to an analog input pin of the microcontroller board.

### Switches

A main power switch was included in the project, and a push button was connected with a pull-down resistance to a digital input pin in order to start the zeroing function.

### Enclosure

A common plastic junction box was used to enclose the micro-controller board and the other electronics; gates were created for the USB connector, the external power supply, the LCD, the main switch, the push button and the connection line to the transducer.

All the electronics components have been purchased at online store.

The wiring diagram of all components was draw using Fritzing, an open source, free software available at http://fritzing.org, and it is showed in Fig.1.

**Figure 1.**
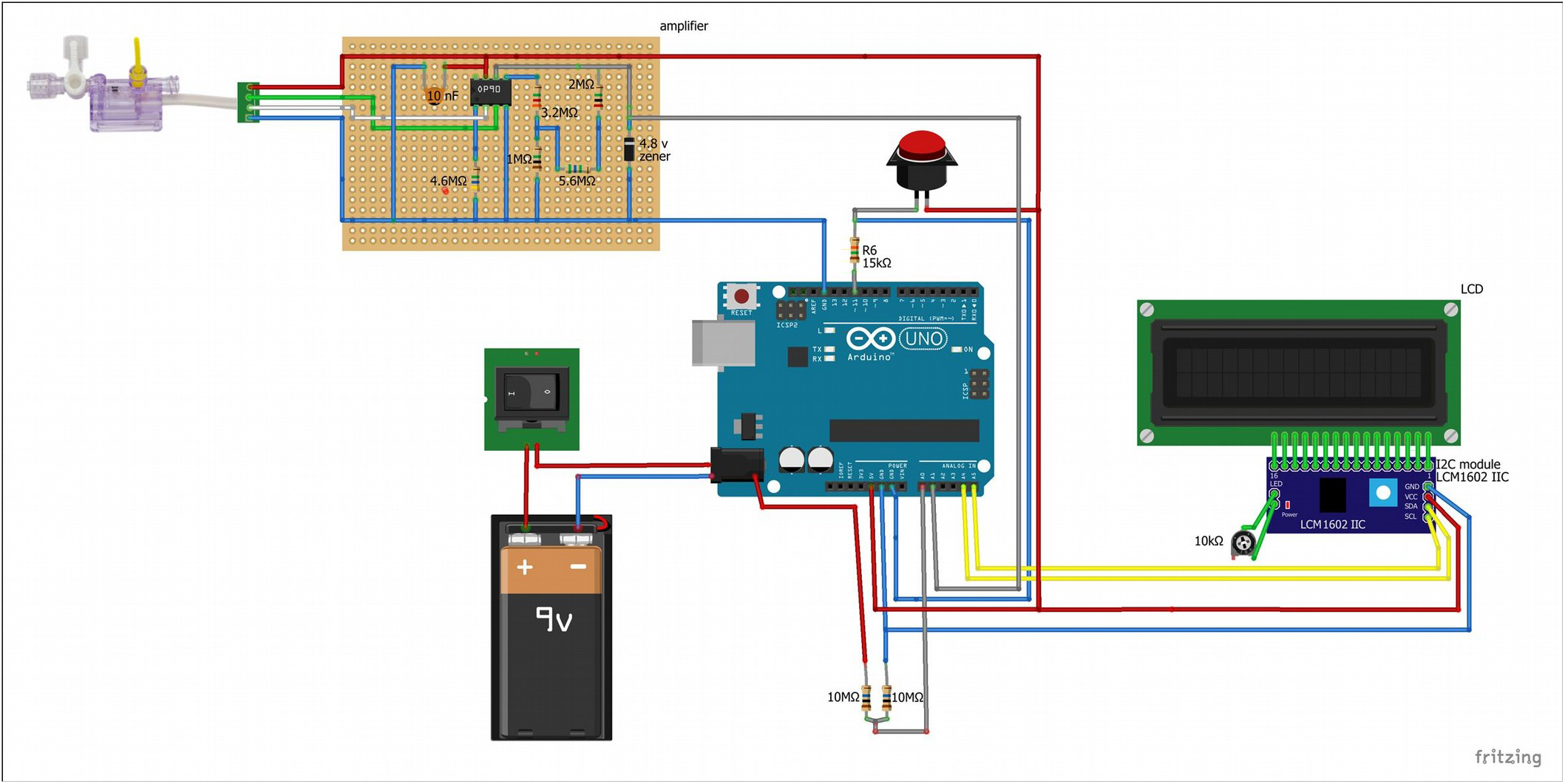
Wiring diagram of the pressure measurement device

### Code

The microcontroller was programmed using the Arduino IDE software version 1.8.1 freely available at the official website [6].

The main program was written in order to cyclically acquire the pressure value from the instrumental amplifier, to calculate a median of 1000 values and to visualize the result on the LCD. During the pressure measurement, a zeroing process can be started by pressing the push button to establish the zero pressure state: a median of 5000 pressure values is calculated, and then subtracted from the following data measurements.

In similar manner, the voltage battery is calculated as the median of 1000 values detected from the voltage divider.

Specific programs were coded to obtain data for the calibration process and to store them in a PC using CoolTerm.

A deboucing function was added into the code to avoid undesired activations of the zeroing process due to transient signals that do not represent a true change of state but can be produced when a switch opens and closes, and the switch contacts bounce off each other before the switch completes the transition to an on or off state.

### Calibration

In this project the calibration was carried out using a water column as standard.

Correlation between pressure and digital input values was obtained with 200 measures for every five cm/H_2_O_2_ from 0 to 35 cm/H_2_O_2_. Conversion from cm/H_2_O_2_ to mmHg was then obtained considering 1 cm/H_2_O_2_= 0,73560 mmHg. Regression line equation was then calculated in order to convert the analog input into pressure measure in mmHg.

A reliability test was performed in another session of 72 pressure measures of water column.

A further test was conducted to compare the device measurements to a standard clinical monitoring system using two Truwave sensors, connected to the same water column.

### Statistics

Correlation test, regression analysis and Bland–Altman analysis were performed using IBM SPSS Statistic v.22.

## Results

The inner and final appearances of the device are shown in Fig.2.

**Figure 2.**
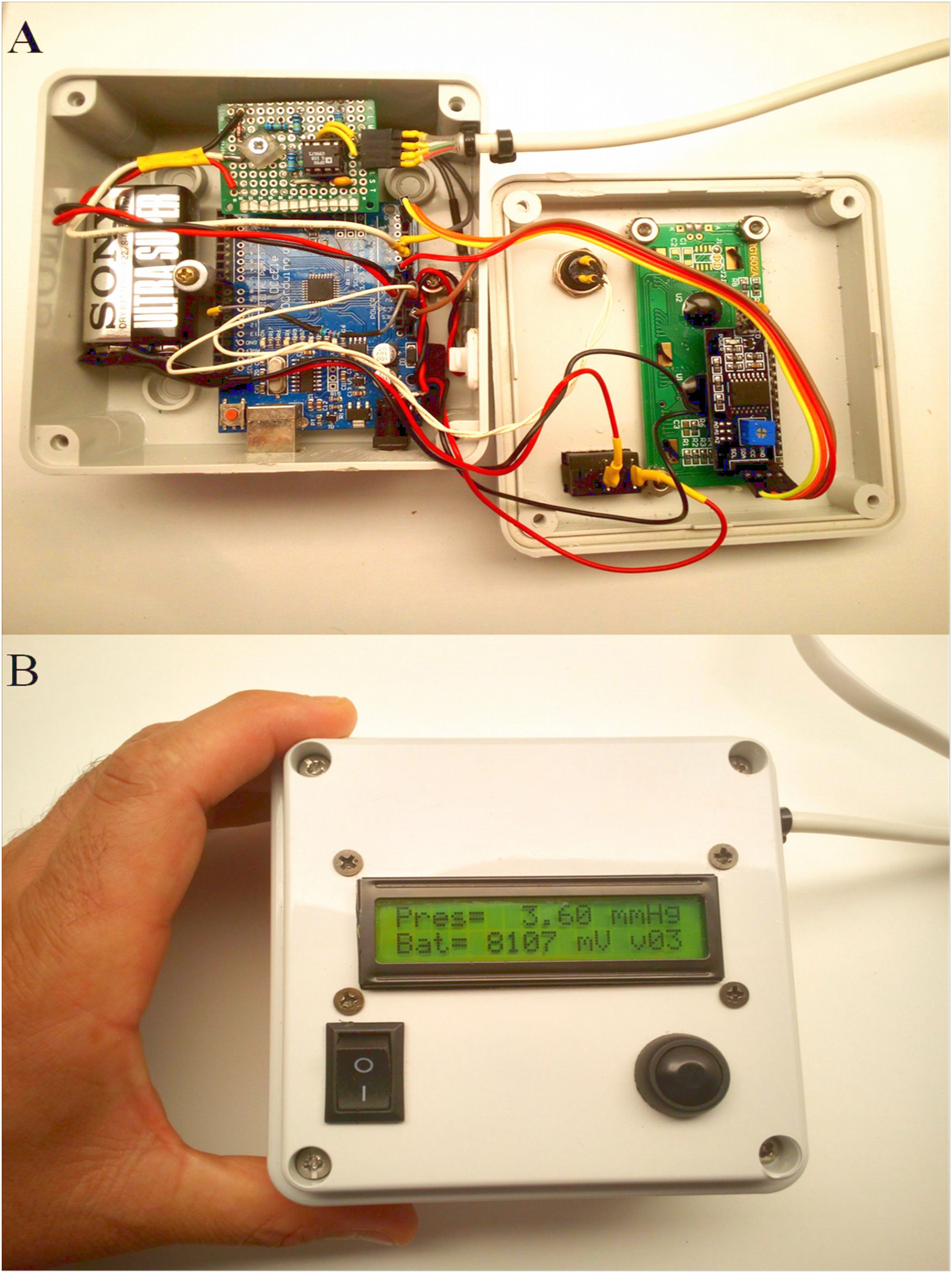
The inner (A) and the external (B) appearance of the device with an LCD showing the pressure value in mmHg in the first row, and the battery charge in mV and the version of the code in the second row. The round push button on the right side starts the zeroing process, while the rectangular one on the left side is the main power switch.

The instrument can be powered using an external power supply (form 6 to 12 V), or using a USB cable connected to a pc, or by the internal 9 V battery: the last option combined with the small size of the plastic box, makes the device handy and portable. The main switch turns on the board in few seconds, while the push button starts the zeroing process that lasts less than two seconds. The LCD output is easily visualized both in dark and bright ambient light due to the trimmable backlight: the pressure values are shown in the first row in mmHg, while in the second row the value of the battery charge expressed in mV and the version of the code are visualized.

The final cost of the device was less than 20 euros (€): microcontroller 4,7 €, integrated circuit OP90 2,5€, LCD and I2C controller 1,86€, plastic junctional box 4€, other electronic components ≈ 5€. Since body fluids are not in contact with the pressure transducers during its use in clinical setting, few of them were recycled for free after accurate cleaning and disinfection with 2% clorexidine.

The calibration process was performed using 1600 pressure measure data: the scattered plot and the correlation coefficient showed a strong linear relationship between the amplified pressure transducer outputs and the reference pressure values; the regression line obtained was therefore use to calibrate our device (Fig.3).

**Figure 3.**
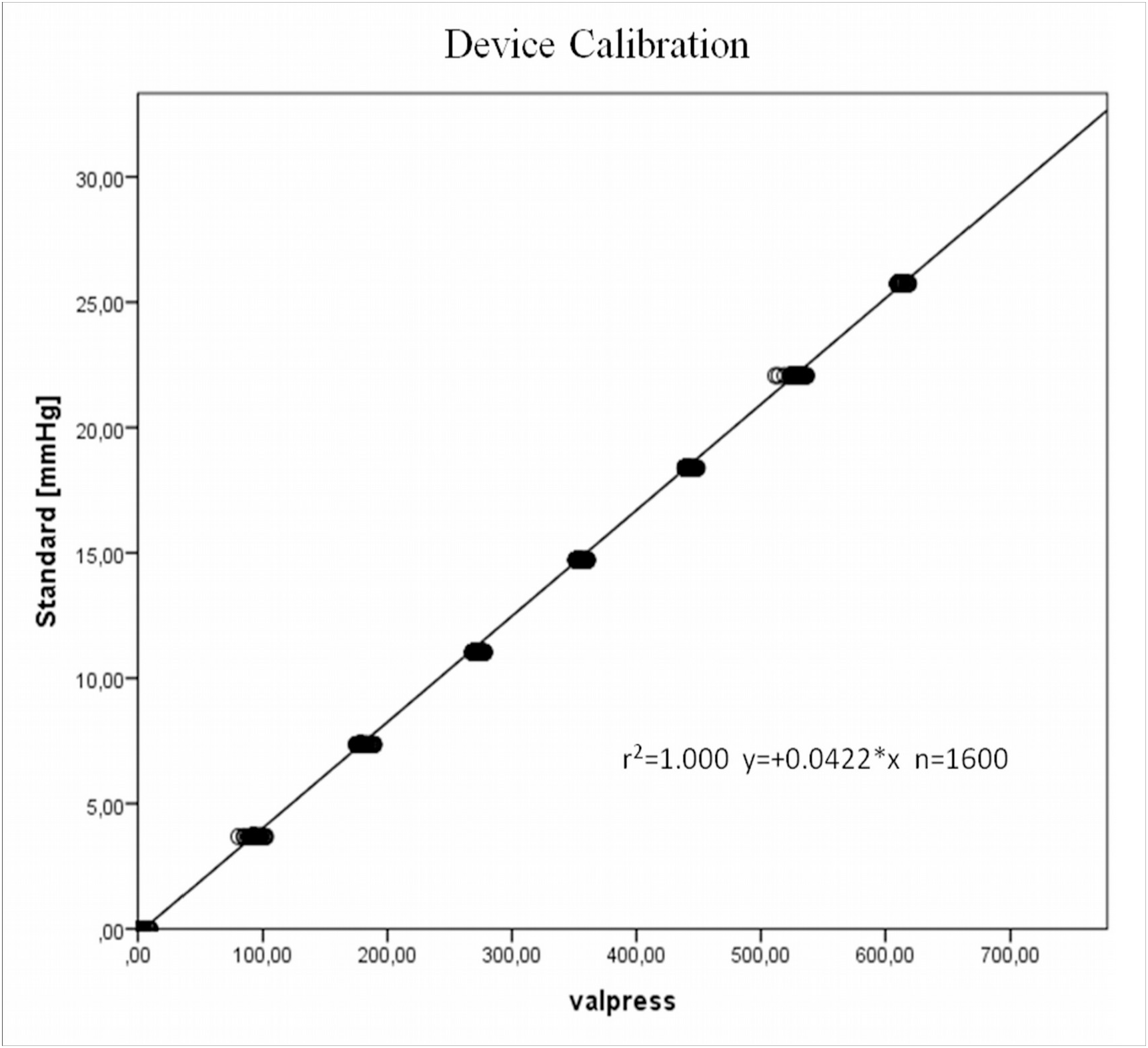
Scattered plot of the 1600 digital value recordered by the Arduino board and the corresponding pressure values in mmHg. R^2^ coefficient shows a strong linear correlation and the regression line is used to calibrate the device.

The consistency of the device was tested with 72 pressure measurements, considering the water column as standard.

The scattered plot and the correlation coefficient showed a strong linear relationship between the measured pressures and the reference pressure values, suggesting agreement between the two different methods of measurement (Fig. 4A).

**Figure 4.**
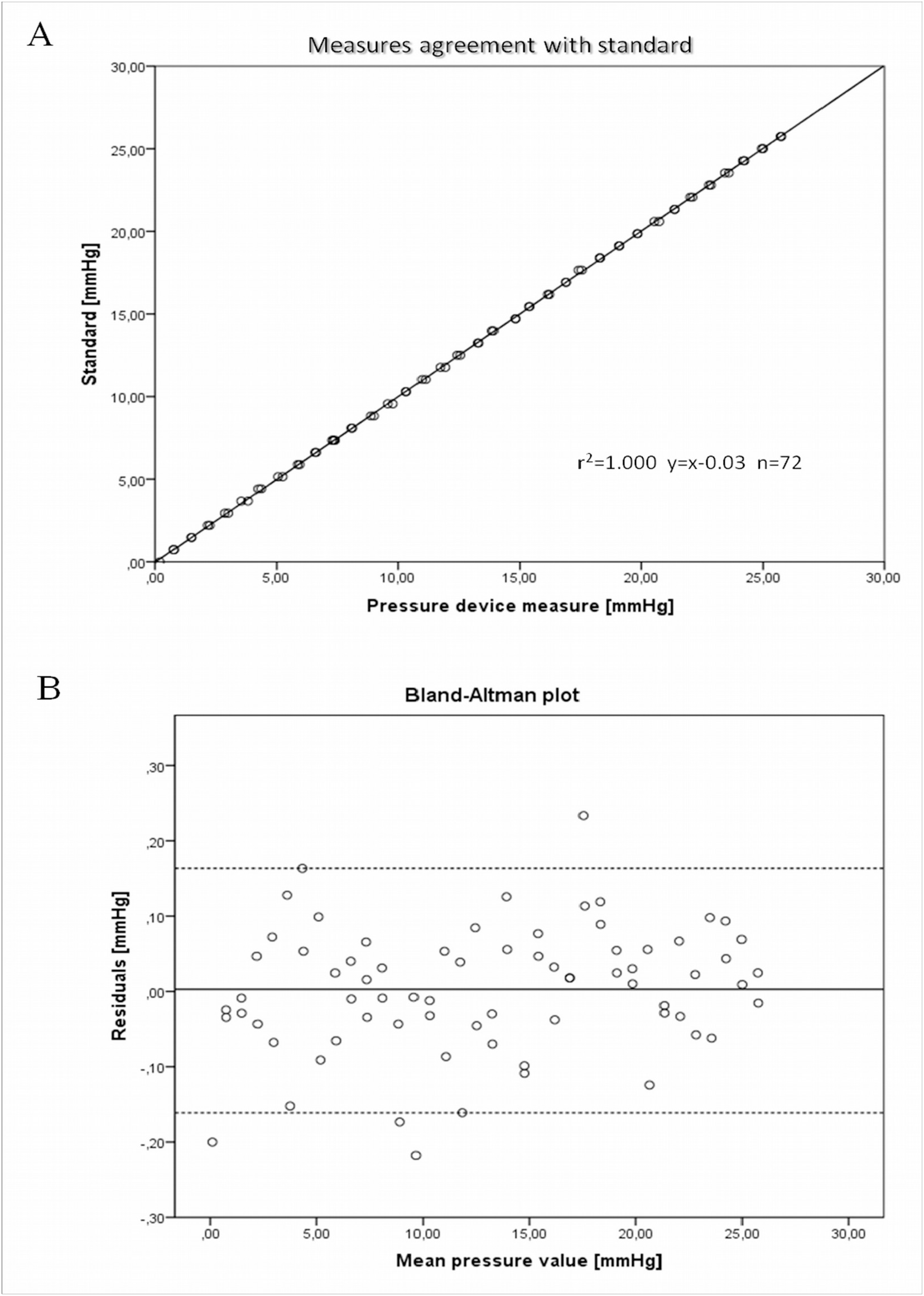
Scattered plot of the 72 pressure values recordered by the device and the corresponding pressure values in mmHg. R^2^ coefficient and regression line show, a strong correlation (A) while the Bland Altman plot (B) suggest the presence of differences between the two methods that are considered not relevant for the purpose of our device.

Residuals were calculated as difference between the standard and the measured values: no significantly statistical difference was found from 0 on the basis of a 1-sample t-test, indicating the absence of fixed bias. A Bland-Altman plot was then created by calculating 95% limits of agreement for each comparison as average difference ± 1.96 standard deviation of the difference: the differences within mean ± 1.96 SD are judged not relevant for the purpose of our device, supporting its use in our research protocol (Fig. 4B).

A further test was performed comparing a standard monitor system used in ICU with our device, both connected to the same water column (Fig.5). Since the resolution of the monitor system was limited to integer values of pressure, the agreement analysis using the pressure measures with decimals from our device was considered not relevant: decimals values were therefore rounded up and compared to the monitor system as standard.

**Figure 5.**
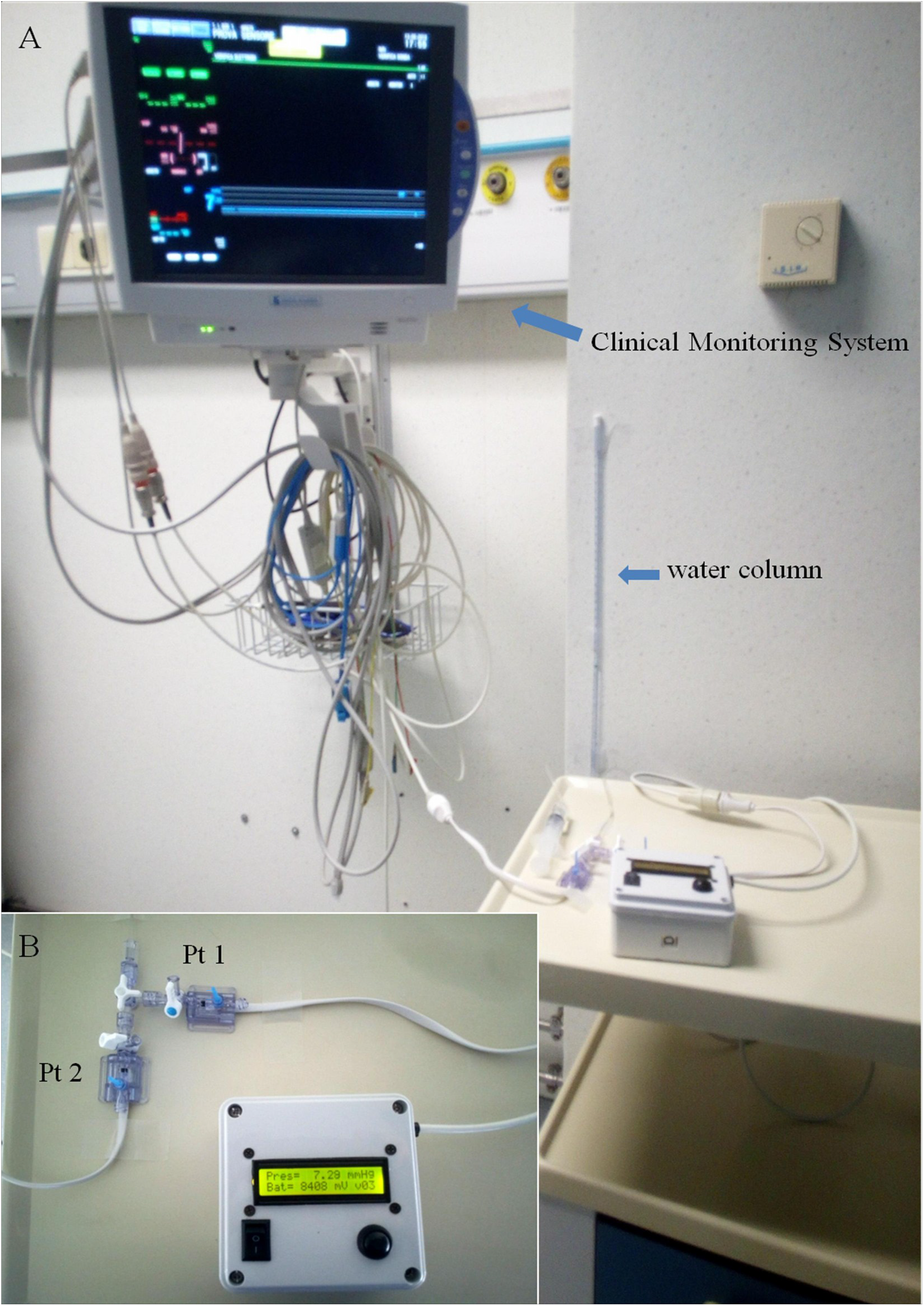
Set-up for measure agreement analysis between a standard clinical monitoring system and the device: the same water column (arrow in A) is connected to two identical pressure transducers Pt1 and Pt2 that are wired to the device (B) and to a standard clinical monitor (arrow in A).

The scattered plot and the correlation coefficient showed a linear relationship (Fig. 6A), while the Bland-Altman plot showed almost no residuals between the two methods (Fig. 6B).

**Figure 6.**
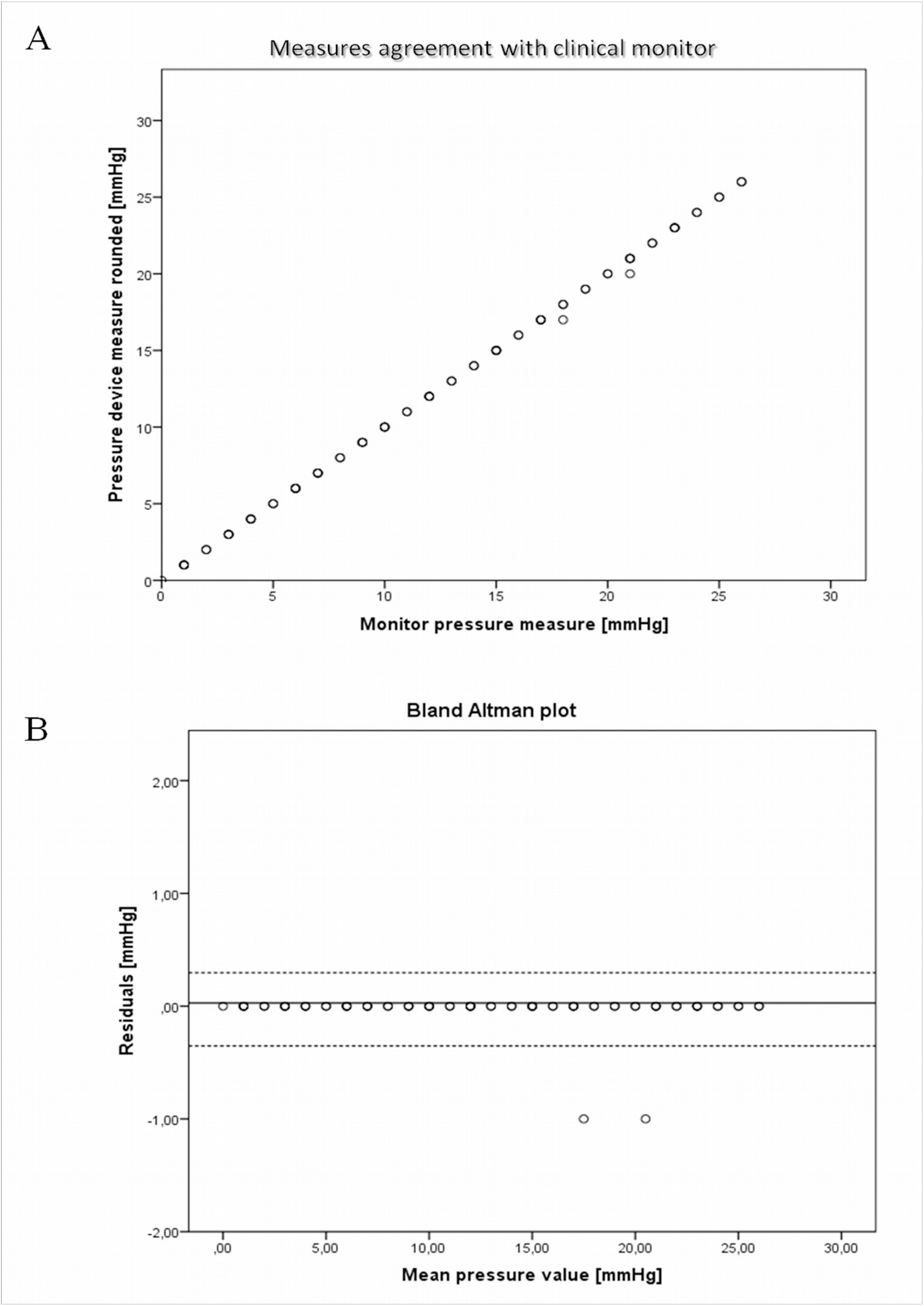
Scattered plot of the 72 pressure values that were rounded reordered by the device and the corresponding pressure values in mmHg recorded by the clinical monitor. R^2^ coefficient and regression line show, an perfect correlation (A) with almost no residuals in the Bland Altman plot (B).

## Discussion

It is a common experience to set experimental procedure using devices already present in lab: usually this process requires simple recognition, collection and arrangement of different instruments; sometimes invasive modifications of the device may be more appropriated for the study purposes, but hacking lab equipment is not straightforward, possible or reasonable; and buying new instruments is not always affordable.

An alternative strategy to obtain new tools is the Do-It-Yourself (DIY) approach; but projecting and building lab instruments seems a tough task limited to electronic and informatics engineers. In 2005 at the Interaction Design Institute in Ivrea (Italy), Arduino platform was created in order to develop a device for controlling interactive-prototypes projected and built by students without a background in electronics and programming. It is an open-source electronics prototyping platform that can be programmed to interact with the world through electronic sensors, lights, and motors. In the last years it has become very popular for several reasons: the hardware is open-source and Arduino-clones are very cheap, the software used to program the board is easy to learn and its free, plenty of sensors and specific shields are available making the board very flexible, and a worldwide community of users has gathered around this open-source platform sharing their knowledge and projects [6,9,10].

The final device herein presented meets the fundamental goals of the original project: it is easy to use and it is portable, it is cheap, both the hardware and the software are easy modifiable, and the pressure measurements are enough accurate to our needs.

However a formal and rigorous calibration and accuracy test was not performed but this device is not aimed to replace industrial ones since any specific lab equipment are probably more accurate. Nevertheless it showed an acceptable accuracy to be employed to test some hypothesis during our research on liver resection in rat.

### Personal learning experience and educational value

Likewise artists, makers and students, many scientists have recognized the potentiality of the free-open-source-hardware (FOSH) movement and some lab tools have been created using Arduino [11, 12], inspiring our project.

I was fascinated by these reports, and I engaged this project with only a previous background in coding BASIC and in electronics experienced at the junior high school, but I was encouraged by curiosity and attitude to the DIY approach [13,14]. It was a valuable learning experience, facilitated by the availability of specific books, on-line video lessons and tutorials, reference websites and by the presence of a large community of makers sharing their experiences.

Indeed, in contrast to the daily work methodology as surgical oncologist, the feeling of being free to fail during the several steps of this project made the acquisition of basic electronics and coding skills really sustainable and the trial and error approach very enjoyable.

The acquisition of basic skills to program the board was a progressive learning process: the final code (Table I) makes the device working as planned, but it could probably be simplified and optimized by more experienced programmers.

**Table I:**
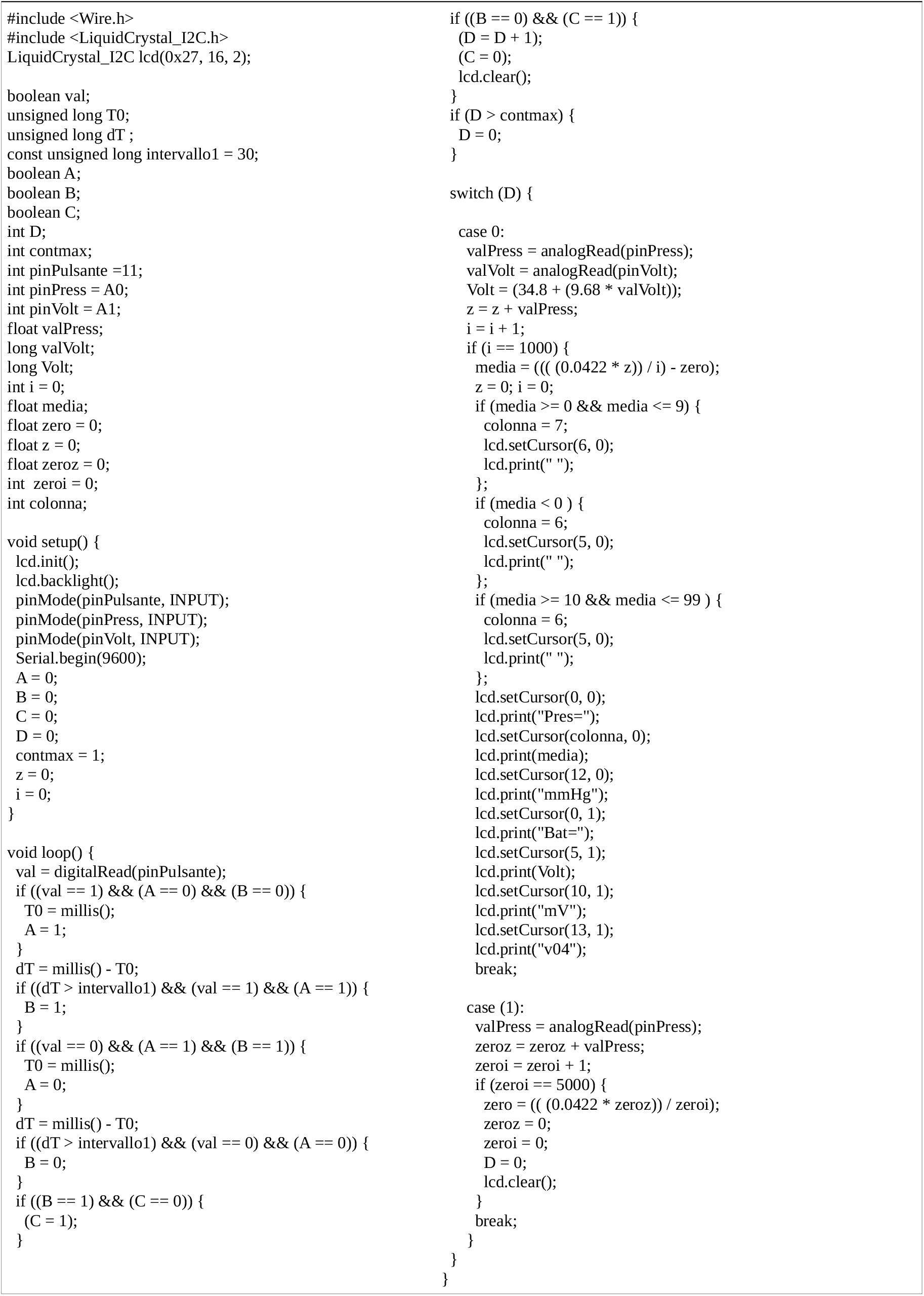
the main code.

The hacking procedures to re-use the sensor transducer and the design and realization of the amplifier could be considered complex and the final result can be obviously optimized. Actually, in my experience, these were the hardest steps of the entire project that however drove me to a deeper understanding of the basics of electronics; since pressure transducers and instrumental amplification boards Arduino-compatible are available, these steps can be easily simplified. Sustained by this preliminary experience and by the interest aroused, a spontaneous working group has been set up by junior colleges and surgical residents, and an Arduino-based platform to monitor vital parameters of small animals and an automated liver perfusion system are presently under development.

## Conclusion

Arduino-board offers an interesting way to improve lab equipment making electronics projects accessible to anyone; it is a valuable learning tool, with which anyone can play and experiment with electronics, learn the foundations and build their own devices.

In a wider contest, Arduino-board and the broader movement of free-open-source-hardware may represent a sustainable and valuable way to enhance scientific education and to increase the opportunities for scientist with limited financial resources.

